# Evaluation of *Octopus maya* enzyme activity of the digestive gland and gastric juice

**DOI:** 10.1101/2024.03.19.585784

**Authors:** Daisy Pineda-Suazo, Wendy Escobedo-Hinojosa, Lenin E. Fabian Canseco, Pedro Gallardo, Cintia Moguel Ojeda, Claudia Caamal-Monsreal, Ariadna Sánchez-Arteaga, Carlos Rosas

## Abstract

**Background:** As the demand for *Octopus maya* grows, sustainable farming practices become essential to prevent overexploitation. Thus, its farming development can be a sustainable alternative to traditional fishing. Understanding the digestive dynamics is essential for devising optimal dietary formulations in aquaculture, particularly the role of enzymes like cathepsins and others. Despite the progress in understanding cephalopod digestion, little is known about the specific functioning of the digestive enzymes responsible for breaking down protein substrates. This knowledge gap underscores the need for further research to ensure *O. maya* population sustainable management.

**Methods and Results:** Dietary formulations are identified for cephalopods by characterizing *O. maya* digestive enzymes present in the digestive gland and gastric juice. The present investigation revealed that acidic proteases showed a peak activity at higher temperatures than alkaline proteases. Inhibitors confirmed the presence of H, L, and D cathepsins. Noteworthy is a lower activation energy of alkaline enzymes compared to acidic, ones highlighting an intriguing aspect of *O. maya’s* digestive physiology.

**Conclusion:** Overall, this research provides valuable insights into *O. maya* digestive enzyme functions representing a significant advancement in formulating diets crucial for octopus successful farming that may help to fully understand its physiology.

## Introduction

*Octopus maya* (*O. maya*) is an endemic species of the Yucatan continental shelf, supporting one of the most important octopus fisheries worldwide with an annual production ranging from 8 000 to 20 000 tons (t) (Markaida et al., 2017). Due to its commercial value, it is one of the top five national fisheries in Mexico (Arreguín-Sánchez et al., 2000). Thus, feeding is a key factor in aquaculture, directly affecting growth, survival, maturation, reproduction, health (Abdollahpour et al., 2022; Vijayaram et al., 2023), and consequently, performance. In the case of cephalopods, our knowledge is still in the early stages of finding feeds with appropriate nutritional composition and satisfactory performance, posing a challenge for large-scale commercial production (Domingues et al., 2007; Rosas et al., 2011).

Research on artificial diet development for cephalopods has focused on understanding nutritional needs regarding total and class lipids, protein content, minerals, and flours. Various raw materials (other cephalopods, fish, and crustacean species); binders (alginate, gelatin); and raw material preparation (fresh, dried or lyophilized) have been recently evaluated for preparing artificial diets (Cerezo Valverde et al., 2008; Estefanell et al., 2013; Garcia et al., 2011; García-Garrido et al., 2010; García-Garrido et al., 2013; Hamdan et al., 2014; Morillo-Velarde et al., 2011; Morillo-Velarde et al., 2012; Morillo-Velarde et al., 2015; Querol et al., 2015a; Querol et al., 2015b; Valverde et al., 2012; Valverde et al., 2013).

Feeds based on natural diets (e.g., crustaceans, fish, and mollusks), wet diets (paste mixtures of natural prey or dried), and formulated diets (agglomerated mixtures of wet and/or dry ingredients with different additives) for growing octopuses have been documented for the common octopus *Octopus vulgaris* (*O. vulgaris*), octopus *O. maya*, monkey-faced octopus *Octopus mimus* (*O. mimus*), and *Enteroctopus megalocyathus* (*E. megalocyathus*) (Gutiérrez et al., 2015; Martínez et al., 2014; Rodríguez-González et al., 2015; Zúñiga et al., 2011).

For octopuses, proteins are the main metabolic substrate characterized by a natural diet primarily based on crustaceans, mollusks, and fish (Alejo-Plata et al., 2018; Estefanell et al., 2013). Previous studies have demonstrated the importance of crustaceans in octopus diets, showing that up to 19 crustaceans species can be found in the diet of *O. vulgaris* wild paralarvae (Roura et al., 2012) and 22 in the diet of *O. maya* juveniles (Markaida, 2023). Thanks to research on physiological digestion, a semi-wet paste based on squid and crab meat was recently developed as a successful diet for *O. maya* juveniles and adults (Gallardo et al., 2020; Martínez et al., 2014; Tercero et al., 2015). With this diet, wild females were successfully acclimated, and juveniles, and pre-adults grew to spawning. The diet was based on the digestive capacity of juvenile octopuses and successfully used to cultivate offspring until they reached 250 g of body weight, as demanded by gourmet market (Rosas et al., 2014).With this diet growth rates of 3.04% per day were reported in juvenile *O. maya* (Martínez et al., 2014). Although, this diet has been well accepted by juveniles, still the challenge is to feed animals with cheaper ingredients as fish scraps from fisheries activities. Until now, diets made with fish scraps are causing lower growth rates than those obtained with squid-crab paste, which has been attributed to lower acceptability (Pérez et al., 2006), conversion index (Domingues et al., 2010), and probably digestibility (Martínez et al., 2014).

To design and develop formulated feed for cephalopods, understanding their digestive physiology is essential. Given the carnivorous feeding habits of octopuses, proteolytic enzymes play a key role in their digestive process; among them are trypsin and chymotrypsin, both from salivary (García-Garrido et al., 2016; Grisley et al., 1996; Morishita et al., 1974; Omedes et al., 2022; Suzumura et al., 2023) and digestive glands, which are the most widely studied enzymes in cephalopods (Mancuso et al., 2014; Martínez et al., 2011; Martínez et al., 2012; Pereda et al., 2009; Rosas et al., 2011). In comparison to proteases, studies on carbohydrases and lipases in cephalopods are limited to *Octopus cyanea* (Boucher-Rodoni, 1973; Omedes et al., 2022), *Eledone cirrosa* (Boucher, 1975), *O. vulgaris* (Boucher-Rodoni and Boucaud-Camou, 1987; O’dor et al., 1984), *Octopus maya* (Aguila et al., 2007; Gallardo et al., 2017; Moguel et al., 2010), *O. bimaculoides* (Ibarra-García et al., 2018; Solorzano et al., 2009) and *O. mimus* (Linares et al., 2015). Therefore, the analysis of digestive enzymatic activity has been considered an indicator of the octopus ability to digest different types of food at different life stages, breeding conditions (Farías et al., 2016) and other environmental factors (Uriarte et al., 2016).

Acidic enzymes in the crop, stomach, and digestive gland (DG) were first observed in *O. vulgaris* (Morishita et al., 1974). Studies by Martínez et al. (2011) and Linares et al. (2015) indicate that acidic enzymes are not only present in *O. vulgaris* but also in *O. maya* and *O. mimus*. These enzymes were also observed in other cephalopod species, such as squid and cuttlefish (Cardenas-Lopez and Haard, 2005; Cardenas-Lopez and Haard, 2009; Perrin et al., 2004), suggesting that they play a key role in the cephalopod digestive capacity. In a partial characterization of *O. maya* digestive enzymes in gastric juice (GJ) and DG (Martínez et al., 2011), cathepsin D is inhibited by 18 and 72% in GJ and DG, respectively, and requires an acidic environment to develop its maximum activity. This situation indicates that acidic enzymes play an important role in the digestive process of this octopus species, as Morishita & Takahashi (Morishita et al., 1974) demonstrated. However, this family of enzymes (cathepsin and pepsin) have shown to be quite sensitive to the biochemical structure of the ingested protein.

To date, the most thoroughly investigated enzymes in cephalopods are those of a proteolytic nature, such as trypsin, chymotrypsin (Boucaud-Camou and Boucher-Rodoni, 1983; Martínez et al., 2011), and cathepsins (Barrett and Kirschke, 1981; Gallardo et al., 2017; Ibarra-García et al., 2018). Given that octopuses are carnivorous organisms with a protein-rich diet (Cerezo-Valverde et al., 2012; Rosas et al., 2013), it follows that the enzymes responsible for protein digestion must be highly efficient and capable of responding rapidly to food arrival in the stomach. Therefore, the hypothesis is that acidic enzymes, such as cathepsins, could initiate and catalyze reactions more efficiently and rapidly compared to alkaline enzymes as chymotrypsin. This phenomenon suggests an evolutionary adaptation by *O. maya*, where acidic enzymes could possess biochemical characteristics that allow them to have lower activation energies, providing a more agile response to the arrival of protein-rich foods in the stomach. This adaptation would align with the requirements of a predatory diet with a high demand for proteins.

The results obtained so far indicate a synchronization between the pulses of DG enzymes and GJ enzymatic activity. In *O. maya*, two pulses were observed (20-80 and 80-180 min), while only one (80-180 min) was observed in *O. mimus* enzymatic activity, suggesting strong differences in digestive dynamics between species (Linares et al., 2015). These differences could be due to the different environmental temperatures in the habitat of each species, with more frequent enzyme release in tropical (e.g*., O. maya*) than in subtropical or temperate species (*O. mimus*). Therefore, temperature could be regulating the entire digestive activity, including ingestion rate, chyme formation, intracellular digestion, and enzyme production.

Given all of the above, exhaustive studies should be performed to fully understand *O. maya* digestive physiology. The relationship between enzymatic activity and the octopus nutritional status, as well as its impact on different life stages and breeding conditions underscores the importance of enzymatic characterization to assess and improve diets. This profound knowledge of digestive physiology will not only contribute to overcoming current obstacles in commercial production but also provide a solid foundation for the sustainable development of the octopus fishery, ensuring the conservation of this species.

## Results

### Determination of the optimal temperature for the activity of acid and alkaline proteases

In the experiments aimed at identifying the optimal temperature for the activity of GJ acid proteases, low activities were noted at temperatures ranging from 10 to 35 °C, with a noticeable surge in the activity observed at 40 and 45 °C (*P* > 0.05; Fig. 1A). In regard to the acid digestive enzymes in the DG, their peak activity showed at 45°C (*P* < 0.05; Fig. 1B). Notably, a rise in activity was also observed from 20 to 30°C, suggesting the potential presence of enzymes with optimum activity within that temperature range (Fig. 1B).

**Fig. 1.**
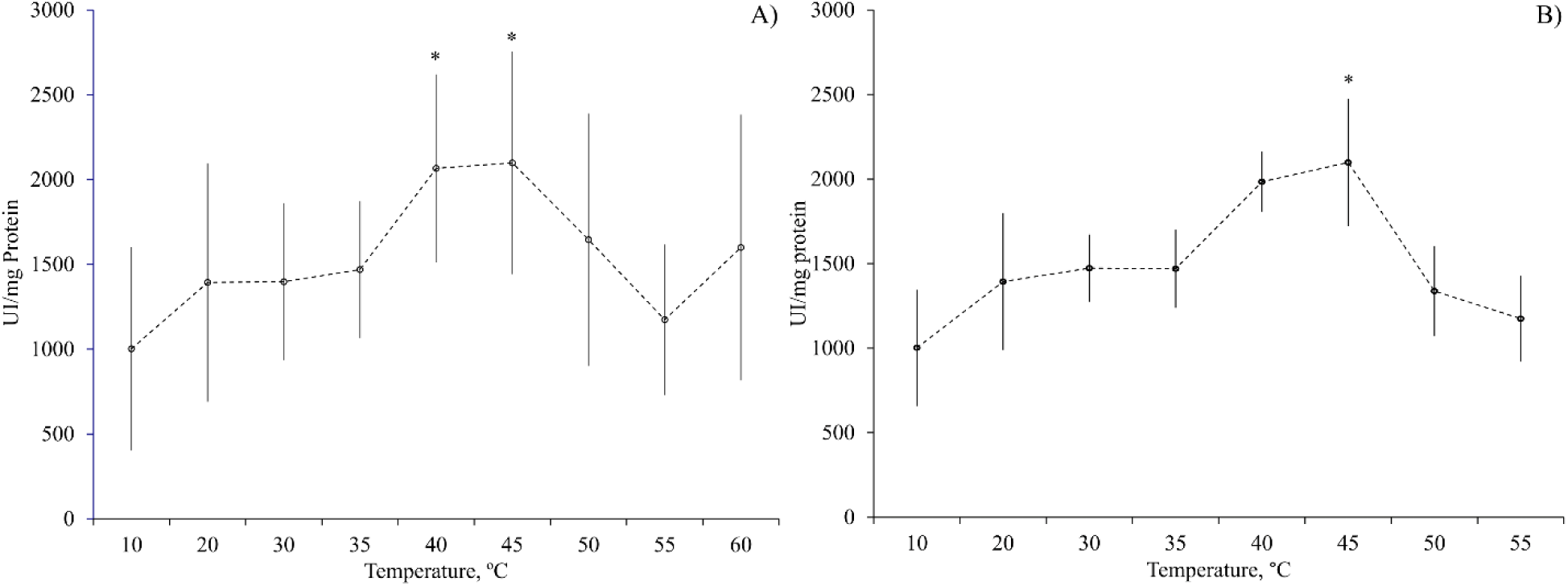
One-way analysis of variance (ANOVA) of temperature dependency on *Octopus maya* acid protease activity. **(A)** Temperature dependency of acid proteases activity of *O. maya* gastric juice; **(B)** Temperature dependency of *O. maya* digestive gland (DG) acid proteases activity. Data show mean ± SE; N = 6, (***P*** < 0.05).

The variance versus average graphs were used to establish which of the temperatures with maximum activity would be chosen as the optimal one for evaluation of acid protease activity of *O. maya* gastric juice. The temperature of 45°C turned out to be the one that promotes the greatest activity. However, at this temperature GJ) enzyme activity turned out to be more dispersed than that observed at 40° C (Fig. 2). Taking the previous into account, the temperature of 40°C was chosen as the optimal one for the evaluation of *O. maya* GJ acidic enzymes.

**Fig. 2.**
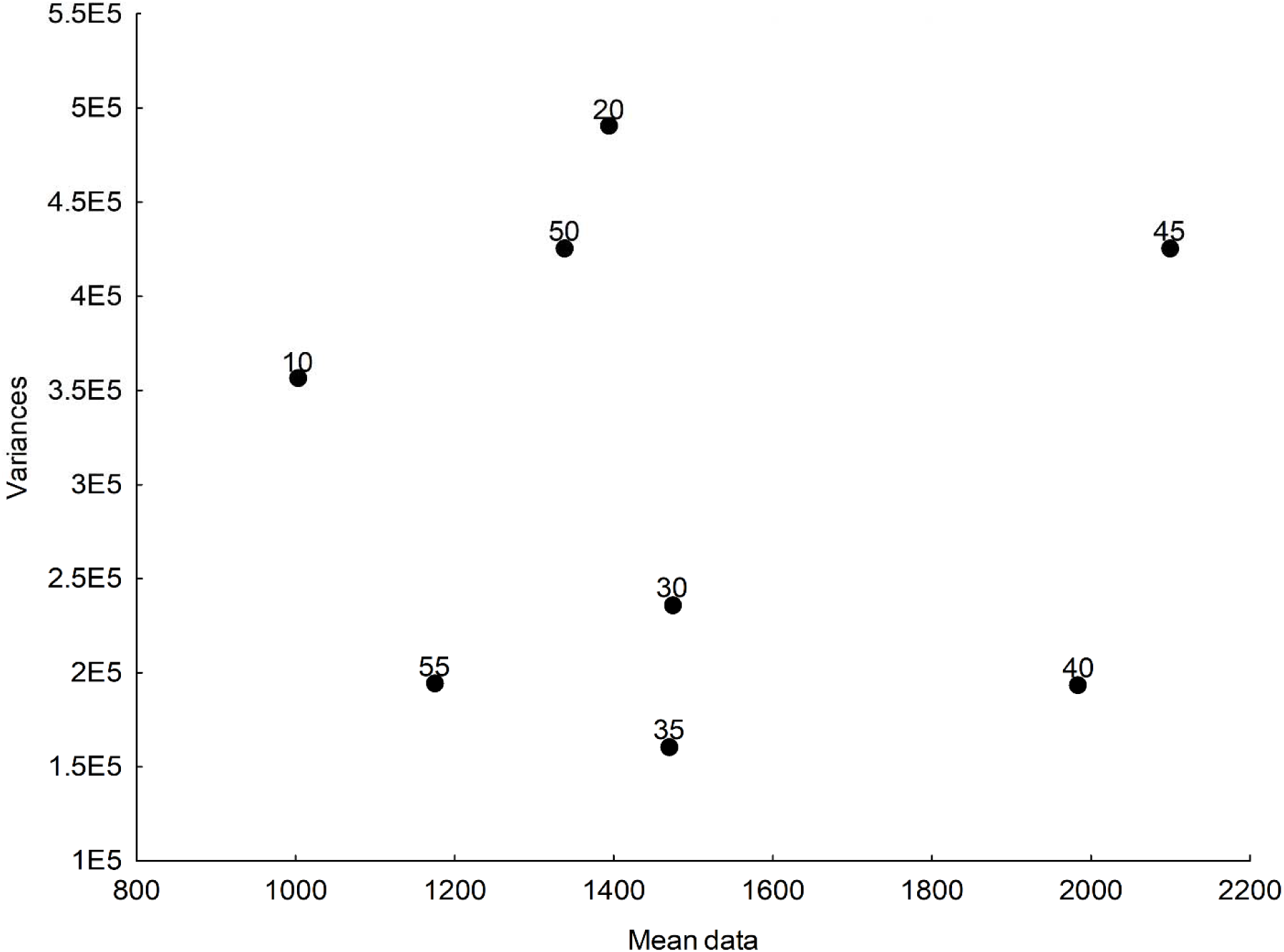
Variance-Average Relationship. Dispersion of variances plotted against the average values of data corresponding to each temperature in the assessment of *Octopus maya* gastric juice (GC) acid protease activity. The x-axis displays the average values of data in U mg protein^-1^, while the y-axis represents the corresponding variances for each average. Each point on the graph represents the average value at each analyzed temperature. The highest protease activity, is observed approximately 1900-2200 U mg protein^-1^ at 40 and 45 °C. However, enzyme activity exhibits greater variability at 40°C.

The GJ alkaline protease enzymatic activity was higher at 35° C (*P* < 0.05; Fig. 3A). The alkaline proteases were observed with relatively lower dispersion at that temperature. The maximum activity of DG alkaline proteases was obtained at 30°C (Fig. 3B). This activity was shown as a single well-defined and highly significant peak (*P* < 0.05).

**Fig. 3.**
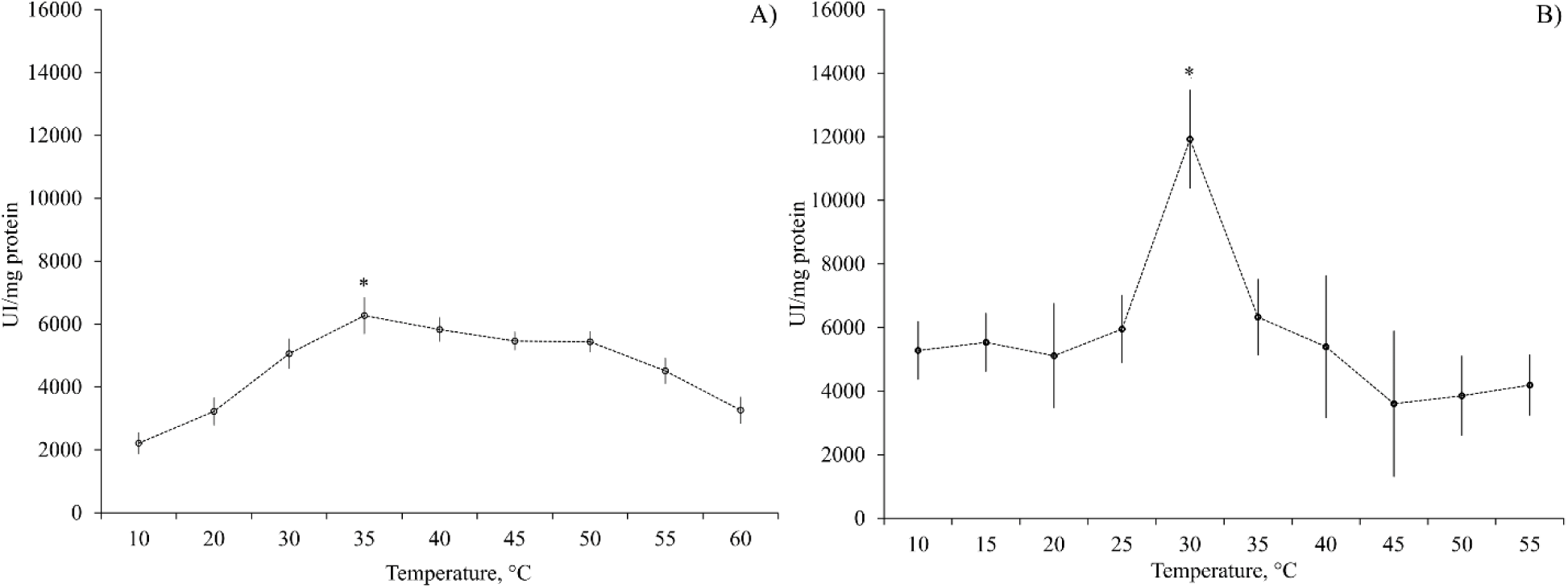
One-way analysis of variance (ANOVA) of *Octopus maya* temperature dependency on alkaline protease activity. **(A)** Temperature effect on *O. maya* gastric juice (**GJ**) alkaline protease activity; **(B)** Temperature effect on *O. maya* digestive gland (**DG**) alkaline protease activity. Data show mean ± standard deviation. N = 6, (*P* < 0.05).

When the enzymatic activity results of alkaline proteases (ranging from 2213 to 11923 U mg protein^-1^) are compared with those of acidic proteases (ranging from 1002 to 2098 U mg protein^-1^), alkaline proteases evidently show notably higher activity levels. Particularly, the DG alkaline proteases displayed the highest activity value recorded, reaching 11293 U mg protein^-1^.

### Activation energy

The activation energy (*Ea*) of all the tests was calculated from the results obtained (Fig. 4), of which the highest *Ea* value was obtained for the peak of maximum activity recorded at 45°C in the acidic proteases from the DG (Fig. 4C), while the lowest one was at 35°C for the alkaline DG proteases (Fig. 4D). Intermediate *Ea* values were obtained for alkaline and acidic proteases of the gastric juice (Fig. 4A and B). Similarly, an *Ea* intermediate value was obtained for the maximum activity of the range from 20 to 30°C of the acidic enzymes in DG for which two values of *Ea* were calculated (Fig. 4C).

**Fig. 4.**
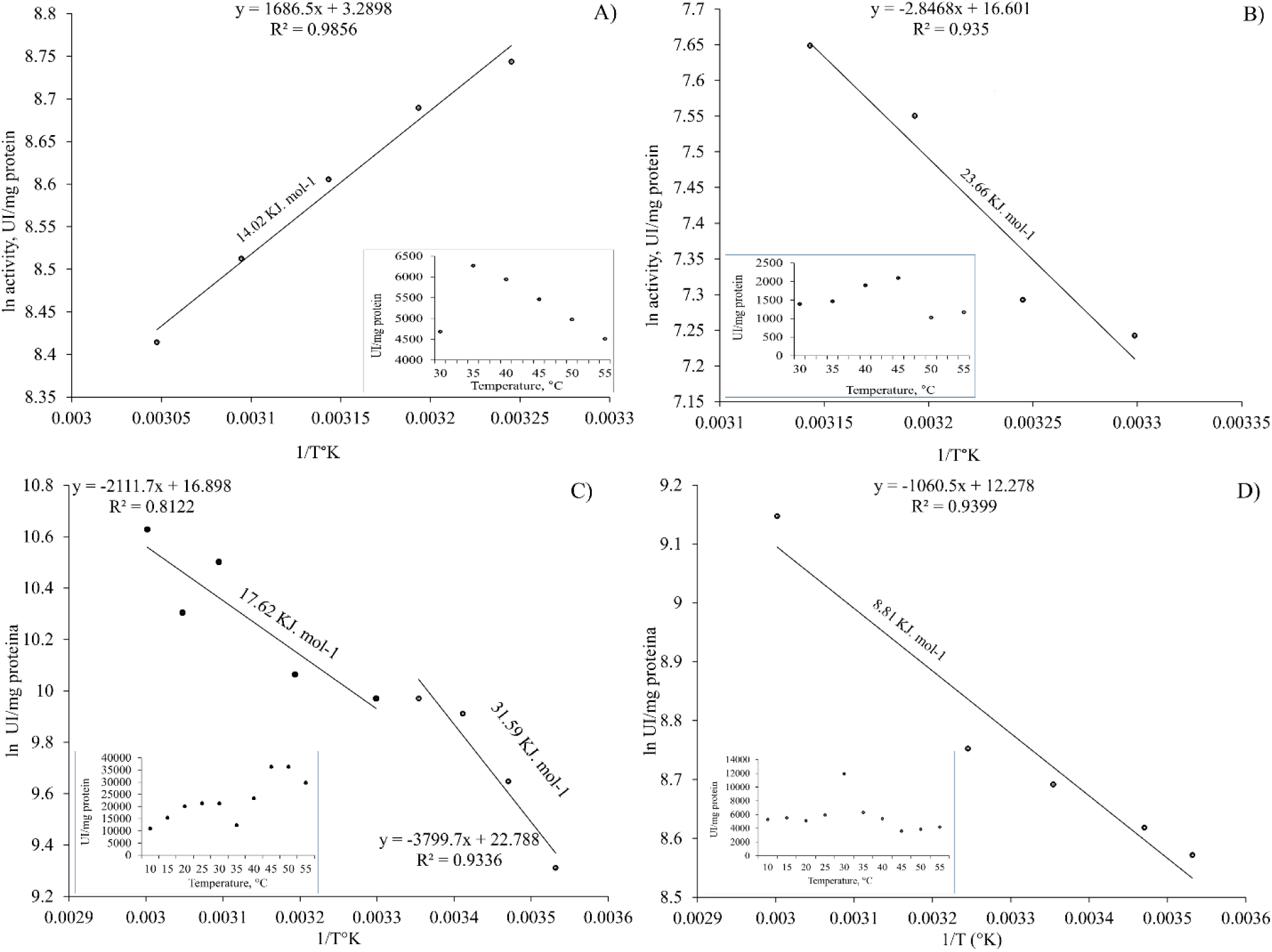
Arrhenius plot analysis for determining activation energy. This figure presents Arrhenius plots utilized to calculate the activation energy of both acidic and alkaline enzymes found in *Octopus maya* gastric juice (**GJ**) and digestive gland (**DG**). The distinct panels (**A**, **B**, **C**, and **D**) delineate the analysis for each enzyme type and tissue location, **A)** Arrhenius plot of the GJ alkaline enzymes; **B)** Arrhenius plot of GC acidic enzymes; **C)** Arrhenius DG acidic enzymes plot; **D)** Arrhenius plot of the alkaline enzymes of DG. The activation energy was calculated using the Arrhenius equation, which relates the reaction rate of a chemical reaction to temperature. The Arrhenius equation is expressed as: 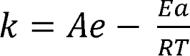 Where: *k* is the reaction rate constant. *A* is the pre-exponential factor or frequency factor, representing the frequency of effective collisions between molecules. *Ea* is the activation energy, representing the minimum energy required for the reaction to occur. *R* is the gas constant (8.314 J/(mol*K)). *T* is the absolute temperature in kelvin. By plotting ln(*k*) versus 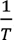, where *k* is the reaction rate and *T* is the temperature in kelvin, the slope of the resulting line on the graph is equal to 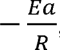, allowing the activation energy *Ea* to be calculated from the slope of the line. These plots elucidate the temperature dependence of enzyme activity and offer insights into the thermodynamic characteristics of enzymatic reactions in *O. maya*. Analysis of temperature dependency is depicted in each Arrhenius plot.

### Determination of the optimal enzyme concentration

Once the optimal temperature was obtained, the tests were carried out to determine the enzyme concentration that maintains a constant activity, observing that from 2 to 16 µL of GJ or DG extract the enzyme activity increased proportionally with the amount of enzyme tested (Fig. 5). The slope values of the relationship between activity and enzyme concentration were 0.0005 (µL GJ) and 0.0002 (µL of DG homogenate) for acidic proteases; 0.0025 (µL GJ) and 7 x 10-5 (µL of DG suspension) for alkaline proteases (Fig. 5). Taking the above into account, a minimum concentration suitable for *O. maya* digestive enzyme evaluation was considered to be 2µL for both the GJ analysis and the DG homogenate.

**Fig. 5.**
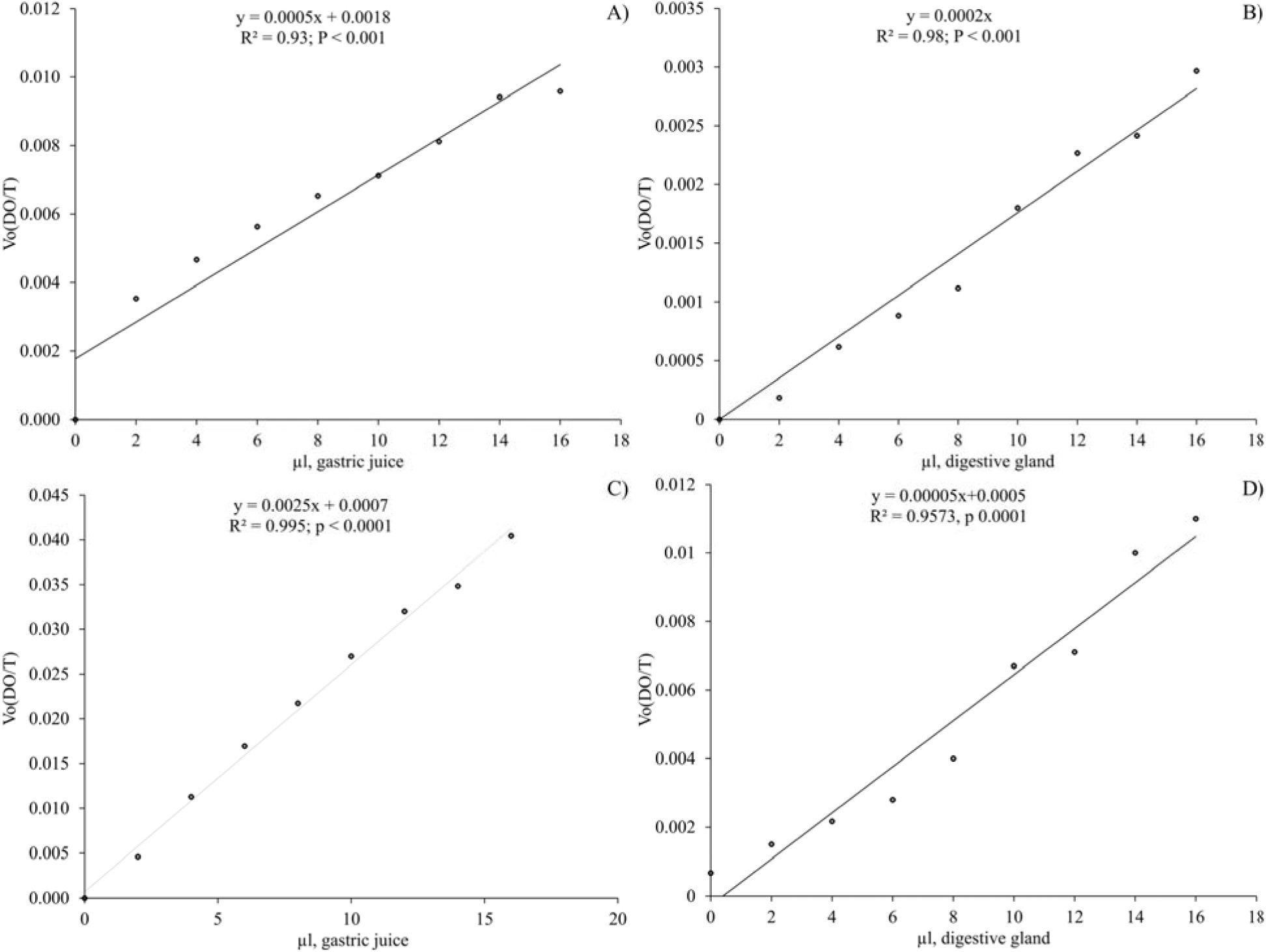
Evaluation of enzymatic variation activity in response to different enzyme concentrations at the identified optimal temperatures. The effect of enzyme concentration from gastric juice (**GJ**) and digestive gland (**DG**) on *O. maya*’s acid and alkaline protease activity, tested at optimal temperatures determined for enzymes from GJ and DG is shown as **A)** Acidic proteases from GJ at 40°C; **B)** Acidic proteases from DG at 45°C; **C)** Alkaline protease from GJ at 35°C; **D)** Alkaline proteases from DG at 30°C. Each graph of linear regression displays R^2^ and P values.

### Effect of inhibitors on cathepsins activity

The inhibition of B, H, L, and D cathepsins was verified by measuring their activities in the presence of E64, leupeptin and pepstatin A inhibitors. Cathepsin H and L activities were significantly reduced by 95.8 and 99.9%, respectively following treatment with E64 inhibitor. Cathepsin L showed a complete enzymatic activity reduction by leupeptin. Cathepsin D did not show enzymatic activity reduction by pepstatin A inhibtor. Cathepsin B showed a 92.5% activity reduction by E64 (Fig. 6).

**Fig. 6.**
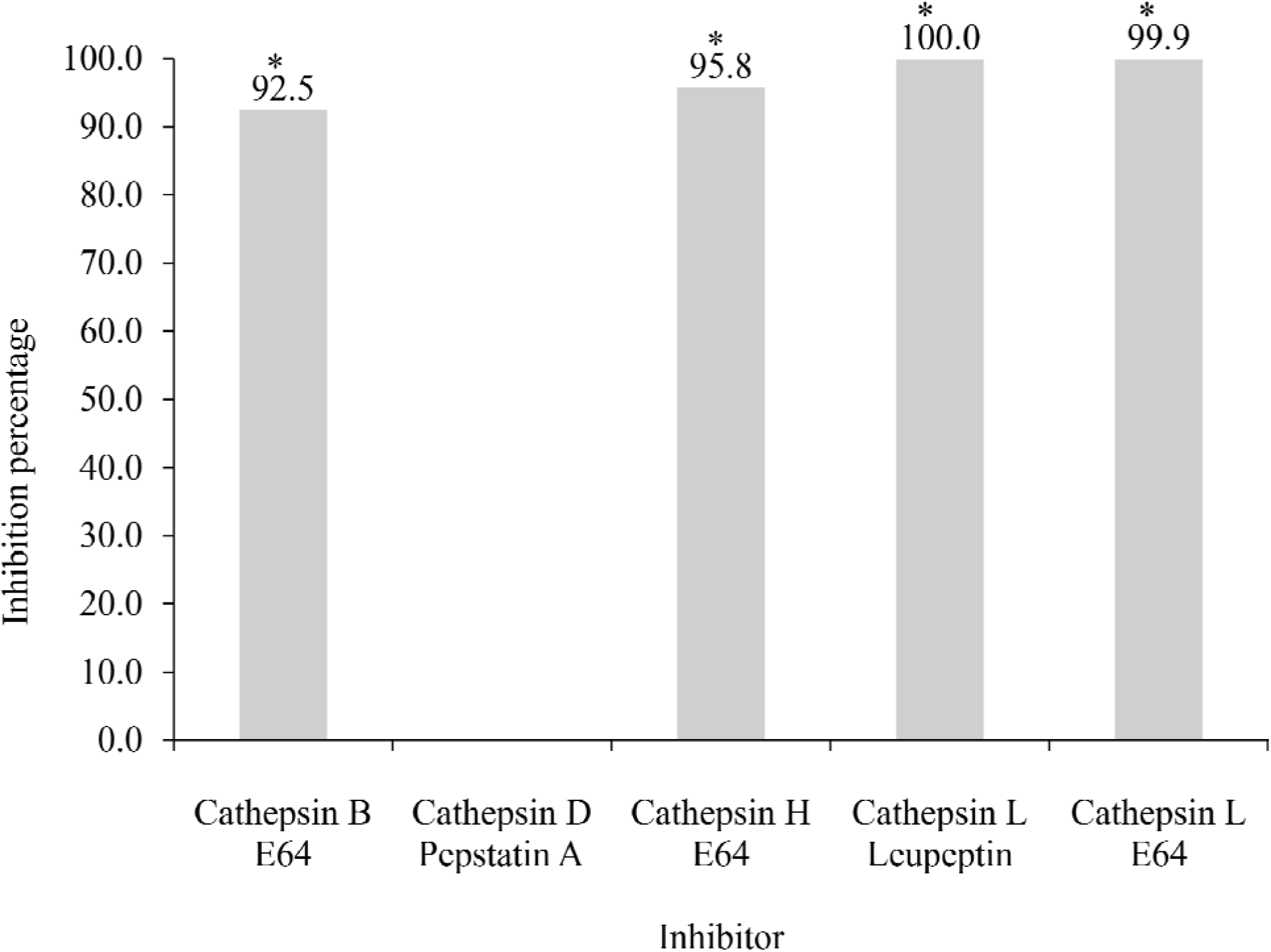
Inhibition percentage by cathepsin enzyme inhibitors present in *Octopus maya* gastric juice (**GJ**): cathepsins **B** by **E64**, **D** by pepstatin **A**, **H** by **E64** and **L** by **E64** and leupeptin inhibitions. Asterisks indicate significant difference (*p* < 0.05). Inhibition percentage = [(**D1** – **D2**)/**D1** × 100] where **D1** = negative control, **D2** = treatment (Pineda-Suazo et al., 2021; Worasatit et al., 1994). The summary of the values used for the inhibition percentage and statistical analysis is given in Table S1.

## Discussion

The digestive anatomy has been extensively characterized in various mollusk species, with a particular focus on cephalopods (Andrews et al., 2022; Omedes et al., 2022; Sykes et al., 2017). It comprises muscle tissue, glands (salivary and digestive), and appendices (Andrews et al., 2022; Omedes et al., 2022). The ingested food, accompanied by saltwater, undergoes mixing with secretions from the salivary glands and is transported to the anterior stomach for the initial digestion stages. According to Andrews et al. (2022), the first part of the digestive system (crop) serves as a storage facility for food boluses, exhibiting significant elasticity and functional characteristics resembling striated muscle. The previous aligns with our present study findings, since the volumes of digestive juice in the anterior stomach were maximal during the initial digestion times, ranging from 1 to 14 mL of juice (Gallardo et al., 2017). In contrast, the cecum appears to be a smaller organ with reduced size, volume, and elasticity compared to the anterior and posterior stomach.

Cardenas-Lopez and Haard (2005) report the presence of D, B, H, and L cathepsins in jumbo squid (*Dosidicus gigas*). These observations were confirmed in the present study (except cathepsin D) by using enzyme inhibitor leupeptin, E64, and pepstatin, thereby providing conclusive evidence for the presence of cathepsins in *O. maya*. digestive juice

Leupeptin is a broad-spectrum inhibitor of serine and cysteine proteases widely utilized in global protein purification workflows, showing inhibitory activity against cA, B, and D athepsins (Aoyagi et al., 1969). On the other hand, E64 is a potent irreversible inhibitor for various cysteine proteases, such as papain, B, H and L cathepsins (Katunuma and Kominami, 1995; Vidal-Albalat and González, 2016). Accordingly, these inhibitors have been widely used to identify and characterize proteases and assess their roles in biological processes (Barrett and Kirschke, 1981; Monteriez et al., 1994). To determine the potential involvement of cysteine-proteinases in *O. maya* digestion, the effects of the proteinase inhibitors leupeptin and E64 were investigated. Our findings revealed that E64 inhibitor reduces enzyme activity up to 99.94%, while leupeptin achieves a reduction of 99.96%. This strong data suggests that cysteine proteases play an essential role in *O. maya* digestive processes.

Enzymes are responsible for controlling many biological processes, catalyzing different reactions using substrates with diverse complexity, size, and chemistry. The single most important property of enzymes is the ability to increase the rates of reactions occurring in living organisms, a property known as catalytic activity. Enzymes are considered to lower the activation energy of a system by making it energetically easier for the transition state to form. Because most enzymes are proteins, their activity is affected by factors that disrupt protein structure, like denaturation, as well as by factors that affect catalysts in general. Factors that denaturate enzymes include temperature, pH as well as well as chemicals like heavy metal and detergents; factors that affect catalysts in general include reactant or substrate concentration and catalyst or enzyme concentration (Copeland, 2013; Robinson, 2015).

The lysosomal cysteine proteinases are generally thought to play an important role in the degradation of intracellular proteins (Barrett and Kirschke, 1981). The L cathepsin has a special place among them since it was found to be the most active lysosomal proteinase in the degradation of various protein substrates, such as azocasein, elastin, or collagen (Barrett and Kirschke, 1981; Kirschke et al., 1982; Mason et al., 1989). Like other lysosomal cysteine proteinases, this enzyme is optimally active in slightly acidic media and unstable under neutral or slightly alkaline conditions (Barrett and Kirschke, 1981). Above pH 6.0 and especially above pH 7.0 the activity is rapidly lost (Mason et al., 1989). Turk et al. (1993) evaluated the effect of temperature (5-37° C) at pH 7.4 in L cathepsin showing an activation energy of 174.7 kJ mol-1 (41.8 kcal mol-1). In 1994, Turk (1994) evaluated cathepsin B in the temperature range of 5-30 °C at pH = 8.0, using 50 mM Hepes buffer, containing 100 mM NaCl and 1 mM EDTA. Under these conditions an activation energy of 183.5 kJ mol-1 was calculated from the Arrhenius plot slope. Both results (L and B cathepsins) are comparable between them. Additionally, both values are also very close to that found for the staphylococcal nuclease alkaline-pH-induced unfolding (195.0 KJ mol^-1^) (Chen et al., 1991). In our experiments, the proteases of the digestive gland proteases were analyzed at pH 6 and 8, covering a range of temperatures from 20 to 45°C. The highest *Ea* value was observed at 45° C (31.59 KJ mol^-1^), intermediate values (17.62 KJ mol^-1^) at 20-30°C and lowest value (8.81 KJ mol^-1^) at 35°C. The variability in *Ea* values among various conditions and enzymes could reflect the complexity of *O. maya* digestive processes. These results suggest the possible presence of digestive enzymes other than B and L cathepsins.

An interesting emerging observation was that alkaline enzymes had a lower activation energy compared to acidic enzymes. This finding led us to question our previous assumptions about the predominant enzymes in *O. maya* digestion, as we had initially assumed that acidic enzymes, such as cathepsins, would have a lower activation energy due to the high-protein carnivorous diet. characteristic of this species. However, it was not the case (Fig. 4), prompting us to reconsider our hypotheses. The proposal is that in addition to cathepsins, alkaline enzymes, such as alkaline phosphatase could play a role in the absorption of amino acids and, in turn, could be involved in cathepsin activation. Possibly, octopuses absorb the enzymes present in their prey at the beginning of external digestion. If these preys (mostly crustaceans) contain alkaline enzymes, such as trypsin, upon entering the digestive tract they could activate the zymogens stored in the anterior stomach, preparing them for the chyme arrival. This activation would be crucial since zymogens remain inactive until food arrives.

Furthermore, alkaline proteases may have evolved to function efficiently in both acidic and alkaline environments, which could be due to pH variability in the gastrointestinal tract that would require the enzymes to be active under different conditions. Therefore, alkaline proteases could have evolved a structure and catalytic mechanism that allows them to have a lower activation energy in an acidic environment as the stomach. This situation could be supported by the results found by Martínez et al., (2011), where activity was observed at pH 7 and 8 in gastric juice, suggesting the presence of alkaline enzymes. When studying the activity of digestive enzymes was studied throughout the digestive tract and over digestion time, Linares et al., (2015), observed alkaline enzyme activity peaks, which were interpreted as changes in pH as digestion progressed. Initially, the pH was acidic, and as acidic enzymes degraded proteins, OH-ions were released, facilitating the activation of alkaline enzymes present in the digestive tract. Then, a shift was observed and the activity of the alkaline enzymes decreased, followed by an increase in the acidic enzyme activity, which was interpreted as the pulse result of enzymes coming from the digestive gland that “boost” digestion by sending “fresh” acidic enzymes to complete it.

Talking about the role of phosphatases as a cofactor to activate proteases, such as trypsin and chymotrypsin, it might be indicative that specific interactions between alkaline proteases and other components present in the stomach could decrease their activation energy. For example, certain cofactors or ions present in the acidic environment of the stomach could stabilize the structure of alkaline proteases and facilitate their catalytic activity, thus reducing their activation energy.

Finally, the activity of proteases, both acidic and alkaline, should be subjected to regulation by other factors, such as post-translational modifications or interactions with other proteins present in the stomach environment. These regulations could lead to a reduction in the alkaline protease activation energy compared to acidic proteases in certain specific contexts. Post-translational modifications have the potential to affect enzymatic activity by modifying the three-dimensional protein structure, which in turn can influence its ability to interact with an affinity for its substrate. These changes could lead to a decrease in the activation energy required to initiate the enzymatic reaction, suggesting that alkaline proteases could be more efficient and active in an acidic environment as the stomach. Therefore, more exhaustive research should help to better understand the set of enzymes involved in *O. maya* digestion.

## Conclusion

Mollusk digestive anatomy – particularly cephalopods like *O. maya* – has been extensively studied, revealing a complex system involving muscle tissue, salivary and digestive glands, and appendices. Our investigation confirmed the presence of various enzymes, including cathepsins B, H, and L, in *O. maya* digestive system. Through the use of enzyme inhibitors like leupeptin, E64, and pepstatin A, the significant role of cysteine proteases in *O. maya* digestion was demonstrated. Notably, we observed that alkaline enzymes were observed to be a lower activation energy compared to acidic enzymes, challenging previous assumptions and suggesting a potential role for alkaline phosphatase in amino acid absorption and cathepsin activation. This finding implies a sophisticated evolutionary adaptation to variable pH conditions in the gastrointestinal tract, supported by enzyme activity observations at different pH levels. Moreover, the presence of specific interactions and regulatory mechanisms involving phosphatases and other proteins in the stomach environment may contribute to the observed differences in activation energy between acidic and alkaline proteases. Overall, further comprehensive research should warrant to fully elucidate the diverse enzyme repertoire involved in *O. maya* digestion and its implications for evolutionary adaptation and ecological dynamics.

## Materials and methods

### Sample collection

A group of *O. maya* 28-wild adults (500 ± 250 g live weight) obtained from the coast of Sisal, and San Felipe, Yucatán, Mexico were kept in acclimation for ten days and fed with fresh crab (*Callinectes* spp), maintained at 24°C and 34 UPS in a recirculatory and aerated seawater system, coupled to 20 and 5 µm cartridge filters. After the acclimation period, gastric juice was obtained from animals fasted for 12 hours, stimulating gastric secretion by placing bags with dead crabs.

These bags allowed the octopuses to be in contact with the prey without having the opportunity to ingest them for a period of 20 min. Subsequently, the animals were sacrificedto extract the gastric juice produced. Therefore, animals were placed in cold sea water maintained at 15°C for 5 minutes and were after wards euthanized by cutting between the eyes into the brain. Subsequently, the mantle was cut longitudinally to locate the stomach, which was clamped at both terminal ends in order to extract the juice accumulated between the distal portion of the cecum and the anterior stomach.

A 3 mL syringe was used to perform the extraction. The digestive gland, which is found occupying the central portion of the mantle just behind the eyes, was also separated and sampled. The digestive gland and gastric juice were frozen immediately by placing them in liquid nitrogen. Subsequently, the samples were stored at -80° C until analysis.

Experimental procedures were approved by the Ethics and Scientific Responsibility Commission of Faculty of Sciences at National Autonomous University of Mexico (CEARC/Bioética/25102021). This commission followed the European directrices used in European Union related with the use of cephalopods as experimental animals. Taking into consideration that until now there is no directrices to Page 12/25 the cephalopods embryos, this study was designed to generate a base line to generate knowledge useful to propose directrices to use embryos as experimental animals.

### Evaluation of the physicochemical and biochemical conditions

#### Protein quantification and enzyme activity assays

Total soluble protein concentration was determined (Bradford, 1976) using serum bovine albumin as standard. Acid and alkaline proteinases were characterized in 96 plate for reader using Anson methods (Anson, 1938). Acid proteinases activity was assayed using hemoglobin (1%) as substrate; alkaline protease activity was assayed using casein (1%) as substrate. Briefly, 20 µL of the enzyme extract (dilution 1:10) was mixed with 500 µL of stauffer buffer; 500 µL of freshly prepared substrate in stauffer buffer at the corresponding pH and incubated at 37°C for 10 min. The reaction was stopped by adding 500 µL of 20% (w/v) trichloroacetic acid (TCA) and cooling for 15 min at 4°C. The precipitated undigested substrate was separated by centrifugation at 13,370 g for 15 min. The supernatant absorbance was measured spectrophotometrically at 280 nm against the substrate without enzyme extract (blank). All determinations were done in triplicates and included blanks, which consisted of stauffer buffer, substrate and TCA without enzyme extracts (Table 1). Blanks were incubated as mentioned earlier and read at each pH. To establish the possible acid enzyme (cathepsin D) presence, the enzymatic extracts were incubated with pepstatin A (1 mM dissolved in dimethylsulfoxide (DMSO) as aspartic proteinase inhibitor at pH 3 (DG) and pH 6 (GJ). This enzyme activity assay was used for all the following experiments.

**Table 1.** Conditions of acid and alkaline protease enzymatic activity reactions present in gastric juice (GJ) and the digestive gland (DG).

| Acid proteases |  | Alkaline proteases |
| --- | --- | --- |
| 20 μL of the extract diluted 1:10. |  |  |
| 500 μL 1% hemoglobin. |  | 500 μL 1% casein. |
| 500 μL stauffer buffer at pH 6.0 (GJ), pH 3 (DG). |  | 500 μL of stauffer buffer at pH 8.0. |
| Incubation for 10 min at 37° C. |  |  |
| Reaction stopped by adding 500 μL of 20% TCA. |  |  |
| Temperature blank with 500 μL of 1% hemoglobin, 500 μL of stauffer buffer at pH 6.0 (GJ), pH 3 (DG) and 20 μL of TCA. |  | Temperature blank with 500 μL of 1% casein, 500 μL of stauffer buffer at pH 8.0 and 20 μL of TCA. |

TCA = trichloroacetic acid

#### Determination of the optimal temperature for the acid and alkaline protease activities

The experimental temperatures in this phase of the present study were set from 10 to 60°C with intervals of 10°C between them. To fine-tune maximum activities, 5°C intervals were also included when necessary.

#### Determination of the optimal enzyme concentration

The experimental enzyme concentrations in the reaction mixture were established between the protein mg contained in a range of 2 to 16 µL of GJ or DG. To compare the results obtained, the experiments were carried out for both, the gastric juice and the digestive gland at 37° C and the previously determined optimal temperatures.

### Evaluation of the activity and identification of cathepsins present in the gastric juice of *O. maya* through the use of specific inhibitors

#### Evaluation of the activity of cathepsins

The standard method (Barrett and Kirschke, 1981) was followed to assess the activity of B, D, H, and L cathepsins in the GJ, which involved using cathepsin-specific substrates, as listed in Table 2.

The reaction mixture was prepared with 50 µL of enzyme extract and 300 µL of buffer/activator (pH 6.0) that was used (88 mM KH2PO4 = 11.96 gr./Lt, 12 mM Na2HPO4 = 1.704 gr./Lt, 1.33 mM EDTA = 0.4950 gr./Lt, 2.7 mM cysteine = 0.3267 gr./Lt). Subsequently, the mixture was pre-incubated at 40° C for 5 min. Then, 10 µL of the substrate in DMSO was added and incubated at 40° C for 10 min. The reaction was stopped with 400 µL of the Fast Garnet GBC preparation in mersalic acid-Brij reagent, allowing it to rest to develop the color for 10 min. The reading was made in a spectrophotometer at 520 nm. The blank contained 50 µL of pyrogen-free water + 300 µL of incubation buffer + 10 µL of substrate + 400 µL of the Fast Garnet GBC preparation in mersalic acid-Brij reagent.

#### Effect of inhibitors on the activities of cathepsins

To identify the cathepsin percentage in the GJ, screenings with the cathepsin specific inhibitors E64, leupeptin and pepstatin A were carried out. The reaction mixture for all assays consisted of 50 µL of GJ and 50 µL of the corresponding inhibitor stock-solution (Table 2).

**Table 2.** Type and concentration of inhibitors used to identify cathepsin activities in gastric juice (GJ) and digestive gland (DG) of *Octopus maya* adults.

| Enzyme | Inhibitor concentration | Substrate |
| --- | --- | --- |
| Cathepsin L | E64 (6 Mm) | ZPAAFC (20 µM) |
|  | Leupeptin (1 µg/mL) |  |
| Cathepsin B | E64 (6 Mm) | BANA (N-Benzoyl-DL-arginineb-naphthylamide) (40 mg/mL) |
|  | Leupeptin (1 µg/mL) |  |
| Cathepsin H | E64 (6 Mm) | H-Arg-AMC (1 µM) |
|  | Leupeptin (1 µg/mL) |  |
| Catepsin D | Pepstatin A (1 µg/mL) | MCA-Gly-Lys-Pro-Ile-Leu-Phe- |
|  |  | Phe-Arg-Leu-Lys(DNP)-DArg- |
|  |  | NH<sub>2</sub> (2.5 µM) |

### Statistical analysis

Standard statistical techniques were used to analyze the data obtained in this research study. Significance was determined by Student’s t-tests, independently by groups to evaluate the differences between the experimental conditions tested with both GJ and DG enzymes. One-way analysis of variance (ANOVA) to evaluate the differences of temperature dependency on *Octopus maya.* Additionally, a linear regression analysis was performed to examine the relationship between enzyme activity and substrates. All statistical analyses were performed using STATISTICA data analysis software (StatSoft, Inc. 2007, version 7. http://www.statsoft.com).

## Supporting information

**Table S1.** Data for inhibition percentage calculation and t-tests of enzyme inhibitors of cathepsins B, D, H, and L. A statistical significance level of *p* < 0.07 was established to evaluate the inhibition percentage by enzymatic inhibitors of cathepsins B, D, H and L, considering the variability of the enzymatic reactions and the relatively low sample size due to ethical restrictions. Inhibition was considered significant if *p* value was less than this threshold.

|  | Cathepsin B |  | Cathepsin D |  | Cathepsin H |  | Cathepsin L |  |  |  |
| --- | --- | --- | --- | --- | --- | --- | --- | --- | --- | --- |
| Inhibitor | E64 | Negative control | Pepstatin A | Negative control | E64 | Negative control | Leupeptin | Negative control | E64 | Negative control |
| S1 | 212.62 | 2401.76 | 3.90 | 4.49 | 0.18 | 3.16 | 8.40734E-05 | 0.14 | 0.00012326 | 0.14 |
| S2 | 281.95 | 3607.56 | 5.84 | 1.80 | 0.41 | 7.94 | 0.000129529 | 0.28 | 0.00015627 | 0.28 |
| S3 | 150.09 | 2631.97 | 3.60 | 4.73 | 0.00 | 2.92 | 3.56706E-05 | 0.16 | 7.8348E-05 | 0.16 |
| S4 |  |  | 5.02 | 5.72 |  |  | 0.000116201 | 0.26 | 0.00012848 | 0.26 |
| S5 |  |  | 4.00 |  |  |  | 4.84518E-05 | 0.17 | 7.0059E-05 | 0.17 |
| Mean | 214.89 | 2880.43 | 4.45 | 4.19 | 0.20 | 4.67 | 8.27851E-05 | 0.20 | 0.00011128 | 0.20 |
| Std. Dev. | 65.96 | 640.15 | 1.22 | 1.67 | 0.20 | 2.83 | 0.00 | 0.06 | 0.00 | 0.06 |
| Inhibition percentage | 92.54 |  | -6.31 |  | 95.76 |  | 99.96 |  | 99.94 |  |
| $p$ value | 0.00 | | 0.28 | | 0.069 | | 0.001 | | 0.001 | |

Where **S** = sample reported in UI/mg protein
Inhibition percentage = $[(D1 - D2)/D1 \times 100]$ where **D1** = control, **D2** = treatment (Pineda-Suazo et al., 2021; Worasatit et al., 1994).

## References

Abdollahpour, H., Falahatkar, B. and Van Der Kraak, G. (2022). Effect of water temperature and food availability on growth performance, sex ratio and gonadal development in juvenile convict cichlid (Amatitlania nigrofasciata). J Therm Biol 107, 103255.

Aguila, J., Cuzon, G., Pascual, C., Domingues, P. M., Gaxiola, G., Sánchez, A., Maldonado, T. and Rosas, C. (2007). The effects of fish hydrolysate (CPSP) level on Octopus maya (Voss and Solis) diet: Digestive enzyme activity, blood metabolites, and energy balance. Aquaculture 273, 641–655.

Alejo-Plata, M. del C., Ahumada-Sempoal, M. A., Guzmán, S. S. L., Herrera-Galindo, J. E. and García-Madrigal, M. del S. (2018). Diet of *Octopus hubbsorum* (Cephalopoda: Octopodidae) from the Coast of Oaxaca, México. Am Malacol Bull 36, 109–118.

Andrews, P. L. R., Ponte, G. and Rosas, C. (2022). Methodological considerations in studying digestive system physiology in octopus: limitations, lacunae and lessons learnt. Front Physiol 13,.

Anson, M. L. (1938). The estimation of pepsin, trypsin, papain, and cathepsin with hemoglobin. J Gen Physiol 22, 79.

Aoyagi, T., Takeuchi, T., Matsuzaki, A., Kawamura, K., Kondo, S., Hamada, M., Maeda, K. and Umezawa, H. (1969). LEUPEPTINS, NEW PROTEASE INHIBITORS FROM ACTINOMYCETES. J Antibiot (Tokyo) 22, 283–286.

Arreguín-Sánchez, F., Solís-Ramírez, M. J. and González de la Rosa, M. E. (2000). Population dynamics and stock assessment for Octopus maya (Cephalopoda:Octopodidae) fishery in the Campeche Bank, Gulf of Mexico. Rev Biol Trop 48, 323–331.

Barrett, A. J. and Kirschke, H. (1981). Cathepsin B, cathepsin H, and cathepsin L. Methods Enzymol 80, 535–561.

Boucaud-Camou, E. and Boucher-Rodoni, R. (1983). Feeding and Digestion in Cephalopods. In The Mollusca, pp. 149–187. Elsevier.

Boucher, R. (1975). Vitesse de digestion chez les cephalopodes eledone cirrosa (lamarck) et illex illecebrosus (lesueur). Cah.Biol.Mar 16, 159–175.

Boucher-Rodoni, R. (1973). Vitesse de Digestion d’Octopus cyanea (Cephalopoda: Octopoda). Mar Biol 18, 237–242.

Boucher-Rodoni, R. and Boucaud-Camou, E. (1987). Fine structure and absorption of ferritin in the digestive organs ofLoligo vulgaris andL. Forbesi (Cephalopoda, Teuthoidea). J Morphol 193, 173–184.

Bradford, M. M. (1976). A rapid and sensitive method for the quantitation of microgram quantities of protein utilizing the principle of protein-dye binding. Anal Biochem 72, 248–254.

Cardenas-Lopez, J. L. and Haard, N. F. (2005). Cysteine proteinase activity in jumbo squid (Dosidicus gigas) hepatopancreas extracts. J Food Biochem 29, 171–186.

Cardenas-Lopez, J. L. and Haard, N. F. (2009). Identification of a cysteine proteinase from Jumbo squid (Dosidicus gigas) hepatopancreas as cathepsin L. Food Chem 112, 442–447.

Cerezo Valverde, J., Hernández, M. D., Aguado-Giménez, F. and García García, B. (2008). Growth, feed efficiency and condition of common octopus (Octopus vulgaris) fed on two formulated moist diets. Aquaculture 275, 266–273.

Cerezo-Valverde, J., Hernández, M. D., García-Garrido, S., Rodríguez, C., Estefanell, J., Gairín, J. I., Rodríguez, C. J., Tomás, A. and García García, B. (2012). Lipid classes from marine species and meals intended for cephalopod feeding. Aquaculture International 20, 71–89.

Chen, H. M., You, J. L., Markin, V. S. and Tsong, T. Y. (1991). Kinetic analysis of the acid and the alkaline unfolded states of staphylococcal nuclease. J Mol Biol 220, 771–778.

Copeland, R. A. (2013). Evaluation of Enzyme Inhibitors in Drug Discovery: A Guide for Medicinal Chemists and Pharmacologists. In (ed. John Wiley and Sons), .

Domingues, P. M., López, N., Muñoz, J. A., Maldonado, T., Gaxiola, G. and Rosas, C. (2007). Effects of a dry pelleted diet on growth and survival of the Yucatan octopus, Octopus maya. Aquac Nutr 13, 273–280.

Domingues, P., Garcia, S., Hachero-Cruzado, I., Lopez, N. and Rosas, C. (2010). The use of alternative prey (crayfish, Procambarus clarki, and hake, Merlucius gayi) to culture Octopus vulgaris (Cuvier 1797). Aquaculture International 18, 487–499.

Estefanell, J., Socorro, J., Izquierdo, M. and Roo, J. (2013). Growth, food intake, protein retention and fatty acid profile in Octopus vulgaris (Cuvier, 1797) fed agglutinated moist diets containing fresh and dry raw materials based on aquaculture by-products. Aquac Res 45, 54–67.

Farías, A., Martínez-Montaño, E., Espinoza, V., Hernández, J., Viana, M. T. and Uriarte, I. (2016). Effect of zooplankton as diet for the early paralarvae of Patagonian red octopus, *Enteroctopus megalocyathus*, grown under controlled environment. Aquac Nutr 22, 1328– 1339.

Gallardo, P., Olivares, A., Martínez-Yáñez, R., Caamal-Monsreal, C., Domingues, P. M., Mascaró, M., Sánchez, A., Pascual, C. and Rosas, C. (2017). Digestive physiology of Octopus maya and O. mimus: Temporality of digestion and assimilation processes. Front Physiol 8,.

Gallardo, P., Villegas, G., Rosas, C., Domingues, P., Pascual, C., Mascaró, M., Sánchez-Arteaga, A., Estefanell, J. and Rodríguez, S. (2020). Effect of different proportions of crab and squid in semi-moist diets for Octopus maya juveniles. Aquaculture 524, 735233.

Garcia, S., Domingues, P., Navarro, J. C., Hachero, I., Garrido, D. and Rosas, C. (2011). Growth, partial energy balance, mantle and digestive gland lipid composition of Octopus vulgaris (Cuvier, 1797) fed with two artificial diets. Aquac Nutr 17, e174–e187.

García-Garrido, S., Hachero-Cruzado, I., Garrido, D., Rosas, C. and Domingues, P. (2010). Lipid composition of the mantle and digestive gland of Octopus vulgaris juveniles (Cuvier, 1797) exposed to prolonged starvation. Aquaculture International 18, 1223–1241.

García-Garrido, S., Hachero-Cruzado, I., Rosas, C. and Domingues, P. (2013). Protein and amino acid composition from the mantle of juvenile O ctopus vulgaris exposed to prolonged starvation. Aquac Res 44, 1741–1751.

García-Garrido, S., Domingues, P., Saworoski, R. and Rosas, C. (2016). Partial characterization of the enzymatic activity in the gastric fluid and digestive gland of Octopus vulgaris. Aquaculture.

Grisley, M. S., Boyle, P. R. and Key, L. N. (1996). Eye puncture as a route of entry for saliva during predation on crabs by the octopus Eledone cirrhosa (Lamarck). J Exp Mar Biol Ecol 202, 225–237.

Gutiérrez, R., Uriarte, I., Yany, G. and Farías, A. (2015). Productive performance of juvenile Patagonian red octopus (Enteroctopus megalocyathus) fed with fresh preys: are relevant the quantity of protein and energy on diets? Aquac Res 46, 64–75.

Hamdan, M., Tomás-Vidal, A., Martínez, S., Cerezo-Valverde, J. and Moyano, F. J. (2014). Development of an in vitro model to assess protein bioavailability in diets for common octopus (Octopus vulgaris). Aquac Res 45, 2048–2056.

Ibarra-García, L. E., Tovar-Ramírez, D., Rosas, C., Campa-Córdova, Á. I. and Mazón-Suástegui, J. M. (2018). Digestive enzymes of the Californian two-spot octopus, Octopus bimaculoides (Pickford and McConnaughey, 1949). Comp Biochem Physiol B Biochem Mol Biol 215, 10–18.

Katunuma, N. and Kominami, E. (1995). Structure, properties, mechanisms, and assays ofcysteine protease inhibitors: Cystatins and E-64 derivatives.pp. 382–397.

Kirschke, H., Kembhavi, A. A., Bohley, P. and Barrett, A. J. (1982). Action of rat liver cathepsin L on collagen and other substrates. Biochemical Journal 201, 367–372.

Linares, M., Caamal-Monsreal, C., Olivares, A., Sánchez, A., Rodríguez, S., Zúñiga, O., Pascual, C., Gallardo, P. and Rosas, C. (2015). Timing of digestion, absorption and assimilation in octopus species from tropical (Octopus maya) and subtropical-temperate (O. mimus) ecosystems. Aquat Biol 24, 127–140.

Mancuso, M., Giordano, D., Genovese, L. maria, Denaro, M. G. and Caruso, G. (2014). Study of digestive enzymes in wild specimens of Sepia officinalis (Linnaeus, 1758) and Octopus vulgaris (Cuvier, 1797). Cah Biol Mar 55, 445–452.

Markaida, U. (2023). Food to go: prey on the web of Octopus maya reveals its diet. Mar Biol 170, 80.

Markaida, U., Méndez-Loeza, I. and Rosales-Raya, M. L. (2017). Seasonal and spatial trends of Mayan octopus, *Octopus maya*, population dynamics from Campeche, Mexico. Journal of the Marine Biological Association of the United Kingdom 97, 1663–1673.

Martínez, R., Sántos, R., Álvarez, A., Cuzón, G., Arena, L., Mascaró, M., Pascual, C. and Rosas, C. (2011). Partial characterization of hepatopancreatic and extracellular digestive proteinases of wild and cultivated Octopus maya. Aquaculture International 19, 445–457.

Martínez, R., Santos, R., Mascaró, M., Canseco, L., Caamal-Monsreal, C. and Rosas, C. (2012). Digestive dynamics during chyme formation of Octopus maya (Mollusca, Cephalopoda). Aquac Res 43, 1119–1126.

Martínez, R., Gallardo, P., Pascual, C., Navarro, J., Sánchez, A., Caamal-Monsreal, C. and Rosas, C. (2014). Growth, survival and physiological condition of Octopus maya when fed a successful formulated diet. Aquaculture 426–427, 310–317.

Mason, R. W., Wilcox, D., Wikstrom, P. and Shaw, E. N. (1989). The identification of active forms of cysteine proteinases in Kirsten-virus-transformed mouse fibroblasts by use of a specific radiolabelled inhibitor. Biochemical Journal 257, 125–129.

Moguel, C., Mascaró, M., Avila-Poveda, O. H., Caamal-Monsreal, C., Sanchez, A., Pascual, C. and Rosas, C. (2010). Morphological, physiological and behavioral changes during post-hatching development of Octopus maya (Mollusca: Cephalopoda) with special focus on the digestive system. Aquat Biol 9, 35–48.

Monteriez, J. P., Kishore, B. K., Maldague, P. and Tulkens, P. M. (1994). Leupeptin and E-64, inhibitors of cysteine proteinases, prevent gentamicin-induced lysosomal phospholipidosis in cultured rat fibroblasts. Toxicol Lett 73, 201–208.

Morillo-Velarde, P. S., Cerezo Valverde, J., Serra Llinares, R. M. and Garcia Garcia, B. (2011). Energetic contribution of carbohydrates during starvation in common octopus (Octopus vulgaris). Journal of Molluscan Studies 77, 318–320.

Morillo-Velarde, P. S., Cerezo Valverde, J., Hernández, M. D., Aguado-Giménez, F. and García García, B. (2012). Growth and digestibility of formulated diets based on dry and freeze-dried ingredients in the common octopus (Octopus vulgaris). Aquaculture 368–369, 139–144.

Morillo-Velarde, P. S., Cerezo Valverde, J. and García-García, B. (2015). Utilization of diets with different fish oil content in common octopus (Octopus vulgaris Cuvier, 1797) and resulting changes in its biochemical composition. Aquac Res 46, 2871–2884.

Morishita, T., Ryuji, U. and Takashi Takahashi (1974). Participation in digestion by proteolytic enzymes of the posterior salivary gland in Octopus . I. Confirmation of the existence of protein digestive enzymes in the posterior salivary gland. Nippon Suisam Gakkaishi (Bull.Jpn.Soc.Sci.Fish.) 40, 601–607.

O’dor, R. K., Mangold, K., Boucher-Rodoni, R., Wells, M. J. and Wells, J. (1984). Nutrient absorption, storage and remobilization in *octopus vulgaris*. Mar Behav Physiol 11, 239–258.

Omedes, S., Andrade, M., Escolar, O., Villanueva, R., Freitas, R. and Solé, M. (2022). B-esterases characterisation in the digestive tract of the common octopus and the European cuttlefish and their in vitro responses to contaminants of environmental concern. Environ Res 210,.

Pereda, S. V., Uriarte, I. and Cabrera, J. C. (2009). Effect of diet and paralarval development on digestive enzyme activity in the cephalopod Robsonella fontaniana. Mar Biol 156, 2121–2128.

Pérez, M. C., López, D. A., Aguila, K. and González, M. L. (2006). Feeding and growth in captivity of the octopus Enteroctopus megalocyathus Gould, 1852. Aquac Res 37, 550–555.

Perrin, A., Le Bihan, E. and Koueta, N. (2004). Experimental study of enriched frozen diet on digestive enzymes and growth of juvenile cuttlefish Sepia officinalis L. (Mollusca Cephalopoda). J Exp Mar Biol Ecol 311, 267–285.

Pineda-Suazo, D., Montero-Vargas, J. M., Ordaz-Ortíz, J. J. and Vázquez-Marrufo, G. (2021). Growth Inhibition of Phytopathogenic Fungi and Oomycetes by Basidiomycete *Irpex lacteus* and Identification of its Antimicrobial Extracellular Metabolites. Pol J Microbiol 70, 131–136.

Querol, P., Gairin, I., Guerao, G., Jover, M. and Tomás, A. (2015a). Growth and feed efficiency of Octopus vulgaris fed on dry pelleted. Aquac Res 46, 1132–1138.

Querol, P., Gairin, I., Guerao, G., Monge, R., Jover, M. and Tomas, A. (2015b). Effect of two extruded diets with different fish and squid meal ratio on growth, digestibility and body composition of Octopus vulgaris (Cuvier, 1797). Aquac Res 46, 2481–2489.

Robinson, P. K. (2015). Enzymes: principles and biotechnological applications. Essays Biochem 59, 1–41.

Rodríguez-González, T., Cerezo Valverde, J., Sykes, A. V. and García García, B. (2015). Performance of raw material thermal treatment on formulated feeds for common octopus (Octopus vulgaris) ongrowing. Aquaculture 442, 37–43.

Rosas, C., Sánchez, A., Pascual, C., Aguila, J., Maldonado, T. and Domingues, P. (2011). Effects of two dietary protein levels on energy balance and digestive capacity of Octopus maya. Aquaculture International 19, 165–180.

Rosas, C., Valero, A., Caamal-Monsreal, C., Uriarte, I., Farias, A., Gallardo, P., Sánchez, A. and Domingues, P. (2013). Effects of dietary protein sources on growth, survival and digestive capacity of Octopus maya juveniles (Mollusca: Cephalopoda). Aquac Res 44, 1029–1044.

Rosas, C., Gallardo, P., Mascaró, M., Caamal-Monsreal, C. and Pascual, C. (2014). Octopus maya. In Cephalopod Culture, pp. 383–396. Dordrecht: Springer Netherlands.

Roura, Á., González, Á. F., Redd, K. and Guerra, Á. (2012). Molecular prey identification in wild Octopus vulgaris paralarvae. Mar Biol 159, 1335–1345.

Solorzano, Y., Viana, M. T., López, L. M., Correa, J. G., True, C. C. and Rosas, C. (2009). Response of newly hatched Octopus bimaculoides fed enriched Artemia salina: Growth performance, ontogeny of the digestive enzyme and tissue amino acid content. Aquaculture 289, 84–90.

Suzumura, Y., Matsubara, K., Morii, S., Abe, M., Gleadall, I. G., Nishikawa, M., Katayama, A., Nishitani, G., Hukushima, T., Yamazaki, T., et al. (2023). Efficacy of octopus feed encased within a collagen membrane. Fisheries Science.

Sykes, A. V., Almansa, E., Cooke, G. M., Ponte, G. and Andrews, P. L. R. (2017). The digestive tract of cephalopods: A neglected topic of relevance to animal welfare in the laboratory and aquaculture. Front Physiol 8, 264006.

Tercero, J. F., Rosas, C., Mascaro, M., Poot, G., Domingues, P., Noreña, E., Caamal-Monsreal, C., Pascual, C., Estefanell, J. and Gallardo, P. (2015). Effects of parental diets supplemented with different lipid sources on Octopus maya embryo and hatching quality. Aquaculture 448, 234–242.

Turk, B., Dolenc, I., Turk, V. and Bieth, J. G. (1993). Kinetics of the pH-induced Inactivation of Human Cathepsin. Biochemistry 32, 375–380.

Turk, B., Dolenc, I., Zerovnik, E., Turk, D. and Turk, V. (1994). Human Cathepsin B Is a Metastable Enzyme Stabilized by Specific Ionic Interactions Associated with the Active Site1”. Biochemistry 33, 14800–14806.

Uriarte, I., Martínez-Montaño, E., Espinoza, V., Rosas, C., Hernández, J. and Farías, A. (2016). Effect of temperature increase on the embryonic development of Patagonian red octopus *Enteroctopus megalocyathus* in controlled culture. Aquac Res 47, 2582–2593.

Valverde, J. C., Hernández, M. D., García-Garrido, S., Rodríguez, C., Estefanell, J., Gairín, J. I., Rodríguez, C. J., Tomás, A. and García, B. G. (2012). Lipid classes from marine species and meals intended for cephalopod feeding. Aquaculture International 20, 71–89.

Valverde, J. C., Martínez-Llorens, S., Vidal, A. T., Jover, M., Rodríguez, C., Estefanell, J., Gairín, J. I., Domingues, P. M., Rodríguez, C. J. and García, B. G. (2013). Amino acids composition and protein quality evaluation of marine species and meals for feed formulations in cephalopods. Aquaculture International 21, 413–433.

Vidal-Albalat, A. and González, F. V. (2016). Natural Products as Cathepsin Inhibitors.pp. 179–213.

Vijayaram, S., Ringø, E., Zuorro, A., van Doan, H. and Sun, Y. (2023). Beneficial roles of nutrients as immunostimulants in aquaculture: A review. Aquac Fish.

Worasatit, N., Sivasithamparam, K., Ghisalberti, E. L. and Rowland, C. (1994). Variation in pyrone production, lytic enzymes and control of rhizoctonia root rot of wheat among single-spore isolates of Trichoderma koningii. Mycol Res 98, 1357–1363.

Zúñiga, O., Paz, A. O. and Torres, I. (2011). Growth evaluation of octopus (Octopus mimus) from northern Chile fed with formulated diets. Lat Am J Aquat Res 39, 584–592.

